# Epigenomic profiling of stem cells within the pilosebaceous unit identifies PRDM16 as a regulator of sebaceous gland homeostasis

**DOI:** 10.1101/2021.04.12.439446

**Authors:** Rizwan Rehimi, Giuliano Crispatzu, Carlos Andrés Chacón-Martínez, Tore Bleckwehl, Giada Mantellato, Gökcen Gözüm, Mathieu Clément-Ziza, Sara A. Wickström, Catherin Niemann, Carien Niessen, Alvaro Rada-Iglesias

## Abstract

The epidermis consists of different compartments such as the hair follicle (HF), sebaceous gland (SG) and interfollicular epidermis (IFE), each containing distinct stem cell (SC) populations. However, with the exception of the SCs residing within the HF bulge, other epidermal SC populations remain less well understood. Here we used an epigenomic strategy that combines H3K27me3 ChIP-seq and RNA-seq profiling to identify major regulators of pilosebaceous unit (PSU) SC located outside the bulge. When applied to the bulk of PSU SC isolated from mouse skin our approach identified both previously known and potentially novel non-bulge PSU SC regulators. Among the latter, we found that PRDM16 was predominantly enriched within the Junctional Zone (JZ), which harbors SC that contribute to renewal of the upper HF and the SG. To investigate PRDM16 function in the PSU SC, we generated an epidermal-specific *Prdm16* Knock-out mouse model (K14-Cre-Prdm16^fl/fl^). Notably, SG homeostasis was disturbed upon loss of PRDM16 resulting in enlarged SGs, and excessive sebum production, resembling some of the features associated with human acne and sebaceous hyperplasia. Importantly, PRDM16 is essential to shut down proliferation in differentiating sebocytes. Overall, our study provides a list of putative novel regulators of PSU SC outside the bulge and identifies PRDM16 as a major regulator of SG homeostasis.

## Introduction

Adult tissue homeostasis requires equilibrium between cell proliferation and differentiation, which is largely dependent on resident SCs that can both self-renew and differentiate into the various cells types found within each tissue (Morrison and Spradling, 2008). SCs reside in unique microenvironments termed “niches”, which are important for the maintenance of the SCs’ unique properties (Fuchs et al., 2004; Watt and Hogan, 2000). Epithelia, one of the four tissue types of the body, are typically highly heterogeneous, containing multiple SC populations that are physically and functionally compartmentalized (Donati et al., 2015). The epidermis is a major example of an adult epithelial tissue in which various SC populations with a spatially confined distribution participate in both skin homeostasis and regeneration upon injury (Jaks et al., 2010; Schepeler et al., 2014). The major components of the mammalian epidermis are the interfollicular epidermis (IFE), the hair follicles (HF) and the sebaceous glands (SG). Several SC populations residing in different locations within the HF and the IFE have been previously described (e.g. HF bulge, HF germ (HG), isthmus, junctional zone (JZ), basal layer of the IFE) (Jaks et al., 2010; Schepeler et al., 2014; Blanpain and Fuchs, 2009). Under normal conditions, each of these SC populations contributes to the homeostasis of specific epidermal compartments. However, upon injury, SCs typically restricted to one compartment can contribute to the regeneration/repair of the whole epidermis (Chuong et al., 2007; Jones et al., 2007; Jensen et al., 2009).

Among the epidermal SCs, the best molecularly characterized population is located within the HF bulge and marked by e.g. keratin 15 and CD34 (Blanpain et al., 2004; Morris et al., 2004). During each hair growth cycle, bulge SC contribute to the formation of the non-permanent lower hair structures (*i*.*e*. hair shaft, inner root sheath, outer root sheath) (Alcolea and Jones, 2014). Furthermore, under homeostatic conditions, bulge SC minimally contribute to the maintenance of the upper part of the HF, the SG or the IFE (Ito et al., 2005). However, upon injury, bulge SC display increased plasticity and differentiation potential, participating in the wound healing response and contributing temporarily to the regeneration of all the other epidermal compartments (Ito et al., 2005). Despite recent advances based on the use of single-cell RNA-seq profiling (Joost et al., 2016; Joost el al., 2018), the SC populations residing within the PSU but located outside the HF bulge (i.e. JZ, SG basal layer, isthmus and infundibulum) remain less well characterized at the molecular level (Jaks et al., 2010; Schepeler et al., 2014). Among them, those SC residing at the intersection between the HF, SG and IFE (*i*.*e*. the JZ) are characterized by expression of high levels of LRIG1. Compared to bulge HFSC, the JZ SC do not express CD34 and show higher proliferative activity (Page et al., 2013). Under homeostatic conditions, these JZ SC maintain the infundibulum and the SG, without contributing to neither the HF nor the IFE (Page et al., 2013). In addition, upon epidermal wounding, JZ SC are mobilized similarly to bulge SC and can participate in the regeneration of all the epidermal compartments (Page et al., 2013). However, in contrast to the bulge HFSC, the intrinsic regulators and transcriptional networks that control the identity and function of JZ SC are not fully understood (Donati et al., 2017; Jensen el al., 2009; Page et al., 2013).

The JZ connects the HF with the SG, an important skin appendage that contains lipid-filled cells known as sebocytes. Sebocytes frequently undergo lysis in order to release a distinct mixture of lipids (*i*.*e*. sebum) that, through a specialized canal, finally reaches the skin surface. Sebum helps to reduce water loss from the skin surface, has antimicrobial activity, is an important source of antioxidants and participates in thermoregulation (Brogden et al., 2012; Fischer et al., 2013; Drake et al., 2008; Schneider and Paus, 2009; Fluhr et al., 2003; Thody and Shuster, 1989). SG homeostasis requires the renewal of the sebocytes undergoing lysis, which depends, at least partly, on Lrig1+ SCs located at the basal layer of the JZ (Jensen el al., 2009; Frances and Niemann 2012; Niemann and Horsley 2012; Thody et al., 1989). Some transcription factors, including PRMD1/BLIMP1 and GATA6, have been implicated in SG maintenance and in controlling the differentiation of Lrig1+ SC into sebocytes (Oules et al., 2019; Oules et al., 2020; Donati et al., 2017; Horsley et al., 2006; Feldman et al., 2019; Kretzchmar et al., 2014). However, the current view of the transcriptional regulators that define the Lrig1+ SC identity and control their contribution to both SG and infundibulum/IFE homeostasis is likely to be incomplete.

We previously showed that, by combining H3K27me3 ChIP-seq and RNA-seq profiling, the main regulators of the different cell types found within a heterogeneous population can be identified by those genes that, although highly expressed, are embedded within broad H3K27me3 domains (Rehimi et al., 2016; Schertel et al., 2015; Shim et al., 2020). Using this simple epigenomic approach to dissect epidermal SC heterogeneity, here we identified several putative regulators of non-bulge SCs. Among them, we found that PRDM16 was co-expressed with LRIG1 within the JZ (Jensen et al., 2009; Frances and Niemann 2012). PRDM16 belongs to the PRDM family of transcription factors and was previously reported as a master regulator of brown fat cell identity (Seale et al., 2008; Kajimura et al., 2009). Here we show that the conditional loss of PRDM16 in the epidermis lead to enlarged and disorganized SGs and impaired sebum secretion. Overall, our work uncovered PRDM16 as a novel marker of the JZ and an important regulator of SG cell fate and homeostasis, which could provide important insights into the molecular basis of common human skin conditions (Shamloul et al., 2020).

## Results

### Epigenomic-based dissection of SC heterogeneity within the pilosebaceous unit

To separate and analyze the bulk of SC present in the PSU but not in the interfollicular epidermis, we used fluorescence activated cell sorting (FACS) to isolate ITGA6^+^/Sca1^-^ cells from the skin of 2^nd^ telogen**-**stage mice (P56) (Fig 1A, Fig 1 – figure supplement 1A). Once ChIP for histone modifications was optimized in these FACS-sorted cells (Fig 1 – figure supplement 1B), RNA-seq and ChIP-seq profiles for H3K4me3 and H3K27me3 were generated (∼5×10^5^ ITGA6^+^/Sca1^-^ cells/ChIP; 1-2 mice) (Fig 1A). The quality of the ChIP-seq profiles is illustrated by the sharp demarcation of the HOX gene clusters into two broad H3K4me3/active and H3K27me3/inactive domains (Fig 1 – figure supplement 1C). Similarly, housekeeping genes with ubiquitous expression (e.g. TBP) were marked by H3K4me3 but not H3K27me3 (Fig 1 – figure supplement 1D), while non-epidermal cell identity regulators (e.g. SOX17) were embedded within broad H3K27me3 domains without H3K4me3 enrichment (Fig 1 – figure supplement 1E).

**Figure 1.**
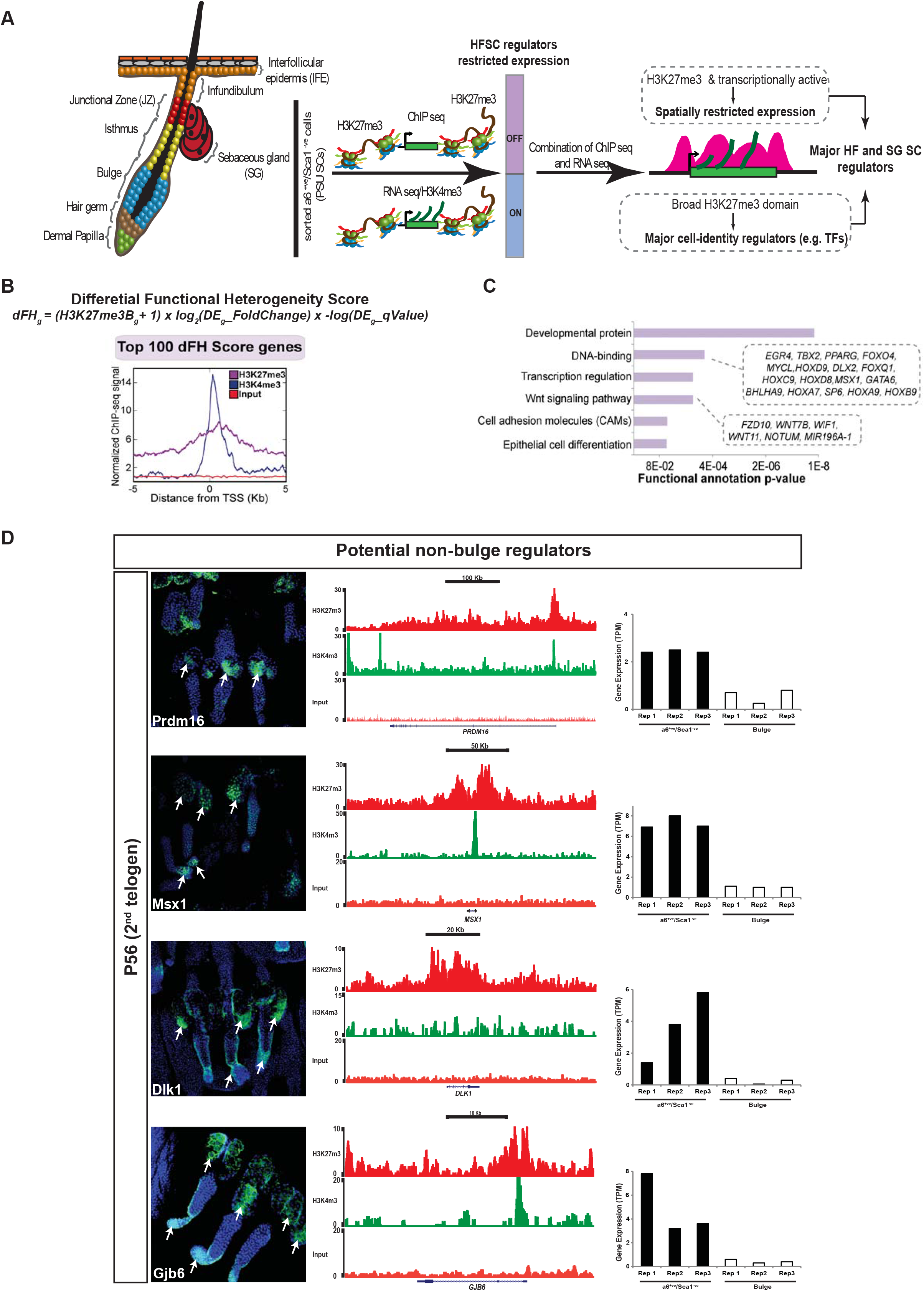
Epigenomic strategy to predict non-bulge stem cell regulators in the mice pilosebaceous unit (PSU). **(A)** Schematic diagram of the epigenomic strategy used to molecularly dissect SC heterogeneity within the *α6^+ve^/Sca1^−ve^* HF and SG cells isolated from the back skin of P56 mice (2^nd^ telogen). Genes with spatially restricted expression within the pilosebaceous unit can be identified as those that are marked by H3K27me3 and transcriptionally active (as measured by RNA-seq). Among them, those covered by particularly broad H3K27me3 domains are likely to represent major SC regulators with important roles in skin homeostasis and/or regeneration. **(B)** For each mouse gene, the dFH score was computed taking into account the size of the H3K27me3 blocks measured in *α6^+ve^/Sca1^−ve^* cells and the differential expression between α6^*+ve*^/*Sca1*^*–ve*^ cells and bulge SC. The average H3K27me3 and H3K4me3 ChIP-seq profiles are shown for the top 100 genes with the highest dFH scores in *α6^+ve^/Sca1^-ve^* cells. **(C)** The top 100 genes with the highest dFH scores in α6^+ve^/Sca1^−ve^ HFSCs were functionally annotated according to Gene Ontology biological process terms. **(D)** ChIP-seq profiles from α6^+ve^/Sca1^-ve^ cells and RNA-seq expression values from α6^+ve^/Sca1^-ve^ cells and bulge SC are shown around *Prdm16, Msx1, Dlk1* and *Gjb6*. Based on immunofluorescence assays the four previous genes showed spatially restricted expression within the pilosebaceous unit that was preferentially observed outside the bulge. All cross-sections were obtained from back skin of P56 (2^nd^ telogen) mice.

To identify important regulators of HFSC populations residing outside the bulge, a differential functional heterogeneity score (dFH score) (Rehimi et al., 2016) was calculated for each gene by multiplying the breadth of the H3K27me3 domain (in bp) around its transcriptional start site (TSS) in ITGA6^+^/Sca1^-^ cells by its differential expression between ITGA6^+^/Sca1^-^ and bulge SC *(dFH=(H3K27_breath+1) x log2FoldChange x -log(differential_expression_qValue)*; Methods) (Fig 1B, Data S1). In principle, genes with high dFH scores should represent major cell-identity regulatory expressed by a subset of PSU SC located outside the bulge (e.g. SG, JZ, Isthmus, hair germ). Next, we ranked all non-bulge genes according to their dFH scores and then focused on those with the highest scores (Fig 1B-C). When analyzed together, the top dFH genes were strongly marked with both H3K27me3 and H3K4me3 and were highly enriched in transcription factors and signaling pathways, some of which were previously implicated in normal skin homeostasis (e.g., *Wnt11, Wif1, Sp6, Fzd10* and *Gata6)* (Fig. 1B-C). Moreover, recently described regulators of non-bulge PSU SC (e.g. *GATA6* and *FOXi3* (Wang et al., 2017; Shirokova et al., 2016; Donati et al., 2017) were also found among the top genes with the highest dFH scores (Fig 1 – figure supplement 1F-G), thus illustrating the specificity and sensitivity of our approach (Rehimi et al., 2016).

### Genes predicted as non-bulge SC regulators display spatially restricted expression within the pilosebaceous unit

We noticed that among the top 200 dFH score genes, there were several that were not previously described as expressed or relevant for PSU SC function. Four of these genes were selected (i.e. *Prdm16, Gjb6, Msx1 and Dlk1*) and their expression patterns were investigated by immunofluorescence in the skin of P56 mice (2^nd^ telogen). Importantly, in agreement with our predictions, all four genes displayed spatially restricted expression within the HF and either low or no expression within the bulge (Fig. 1D). More specifically, PRDM16 was highly expressed within the JZ and the SG, while the other three genes (i.e. MSX1, DLK1 and GJB6) were mostly expressed in the HG and the SG. Moreover, we also observed the expression of DLK1 in lower bulge SC.

Next, we selected *Prdm16* for further functional characterization, as it belongs to the PRDM family of transcription factors, which have major roles in the control of cellular identity under both physiological and pathological conditions (Fog et al., 2012; Hohenauer and Moore 2012). PRDM16 controls the cell fate between muscle and brown fat cells and its loss from brown fat promotes muscle differentiation (Seale et al., 2008). However, its functional relevance during skin homeostasis was unknown. To more precisely define the SC population in which PRDM6 is expressed within the pilosebaceous unit, we performed double immunostainings for PRDM16 and LRIG1 in back skin sections and tail whole mounts. Remarkably, PRDM16 and LRIG1 showed highly overlapping expression patterns during both *2^nd^* telogen (P56) and 1^st^ anagen (P33, especially in the JZ (Fig 2A-B). Moreover, in the tail skin of P56 mice, both proteins were also co-expressed in the HG and the SGs (Fig 2A), although HG expression might be artifactual as it was not reproducibly observed among replicates (Fig 1D). Overall, these results show that PRDM16 represents a novel marker for the JZ SC (*i*.*e*. Lrig1^+^ SC) (Fig 2C) and suggest a potential role in the homeostasis of the SG and/or the infundibulum (Jensen et al., 2009).

**Figure 2.**
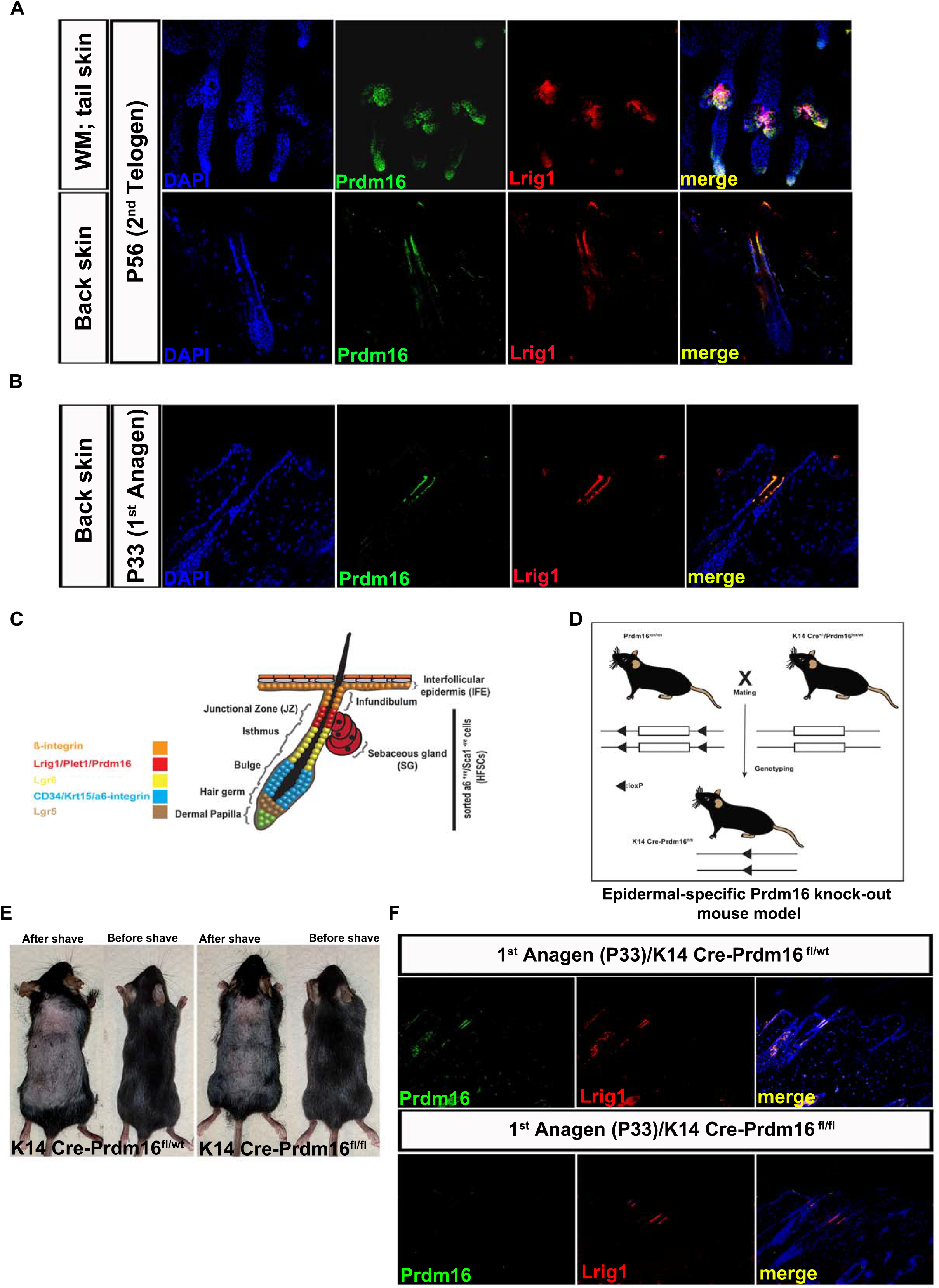
Characterization of PRDM16 expression in the skin and generation of an epidermal-specific Prdm16 Knock-out mouse model. **(A-B)** Double immunofluorescence for PRDM16/LRIG1 in tail skin whole mounts from **(A)** P56 (2^nd^ telogen) mice (upper row) and back skin sections from P56 (2^nd^ telogen) (bottom row) or (B) P33 (1^st^ anagen) mice. PRDM16 extensively co-localizes with LRIG1, particularly within the JZ. **(C)** Schematic diagram of epidermal SC compartments in mouse skin during 2^nd^ telogen (P56). The expression of several SC markers is shown in different colors, with the expression domain of PRDM16 within the JZ showed in Red. **(D)** Schematic diagram depicting the breeding strategy used to generate an epidermal-specific *Prdm16* Knock-out mouse model (K14-Cre-Prdm16^fl/fl^). **(E)** The specific loss of Prdm16 in the epidermis (K14-Cre-Prdm16^fl/fl^) did not result in any obvious morphological or fur differences in comparison with wild-type mice (K14 Cre-Prdm16^fl/wt^). **(F)** Double immunofluorescence for PRDM16 and LRIG1 in back skin sections from P33 (1^st^ anagen) mice that were either WT (K14 Cre-Prdm16^fl/wt^; upper row) or deficient for PRDM16 in the epidermis (K14-Cre-Prdm16^fl/fl^).

### Skin-specific loss of PRDM16 causes enlarged sebaceous glands and increased sebum production

To study the functional role of *Prdm16* during skin homeostasis, we crossed previously generated *Prdm16^fl/fl^* mice (Corrigan et al., 2018) with mice expressing Cre recombinase under the control of the human *Keratin-14* promoter (Hafner et al., 2004) (Fig 2D). This allowed us to obtain K14-Cre-*Prdm16^fl/fl^* mice in which *Prdm16* was specifically deleted in the basal layer of the skin and the PSU, including the undifferentiated cells of the sebaceous gland (Fig. 2D, Fig 2 – figure supplement 1A-B). Upon initial evaluation, the K14-Cre-*Prdm16^fl/fl^* mice did not show any gross morphological abnormalities and their fur appear normal (Fig. 2E). Immunofluorescence analyses in tail skin whole mounts and back skin sections confirmed that PRDM16 was barely detectable in K14-Cre-*Prdm16^fl/fl^* mice, while Lrig1 levels appeared unaffected in both the JZ and the SGs (Fig. 2F and Fig 2 – figure supplement 1C). These results indicate that Lrig1 expression is not affected by the loss of PRDM16.

Next, to evaluate whether any structural changes occurred in the skin of K14-Cre-*Prdm16^fl/fl^* mice, back skin sections from 1^st^ and 2^nd^ telogen (P23 and P56 respectively) mice were stained with H&E. Notably, these stainings revealed that the loss of *Prdm16* led to a clear enlargement of the SGs (Fig. 3A), while the infundibulum and the rest of the HF appeared to be normal. Furthermore, Nile Red staining during multiple stages of the hair cycle (i.e. P23/P33/P56/P63) confirmed that the SGs of the K14-Cre-*Prdm16^fl/fl^*mice were significantly larger than those of the control mice (Fig 3B-D; Fig3 – figure supplement 1A-B). Moreover, in the absence of PRDM16, the SGs seemed to produce an excess of sebum that accumulated in droplets surrounding the SGs (Fig 3B-C; Fig 3– figure supplement 1A-B). This increase in sebum production was also observed upon Oil Red O staining, which showed accumulation of lipids in the IFE and dermis of K14-Cre-*Prdm16^fl/fl^* mice (Fig 3E).

**Figure 3.**
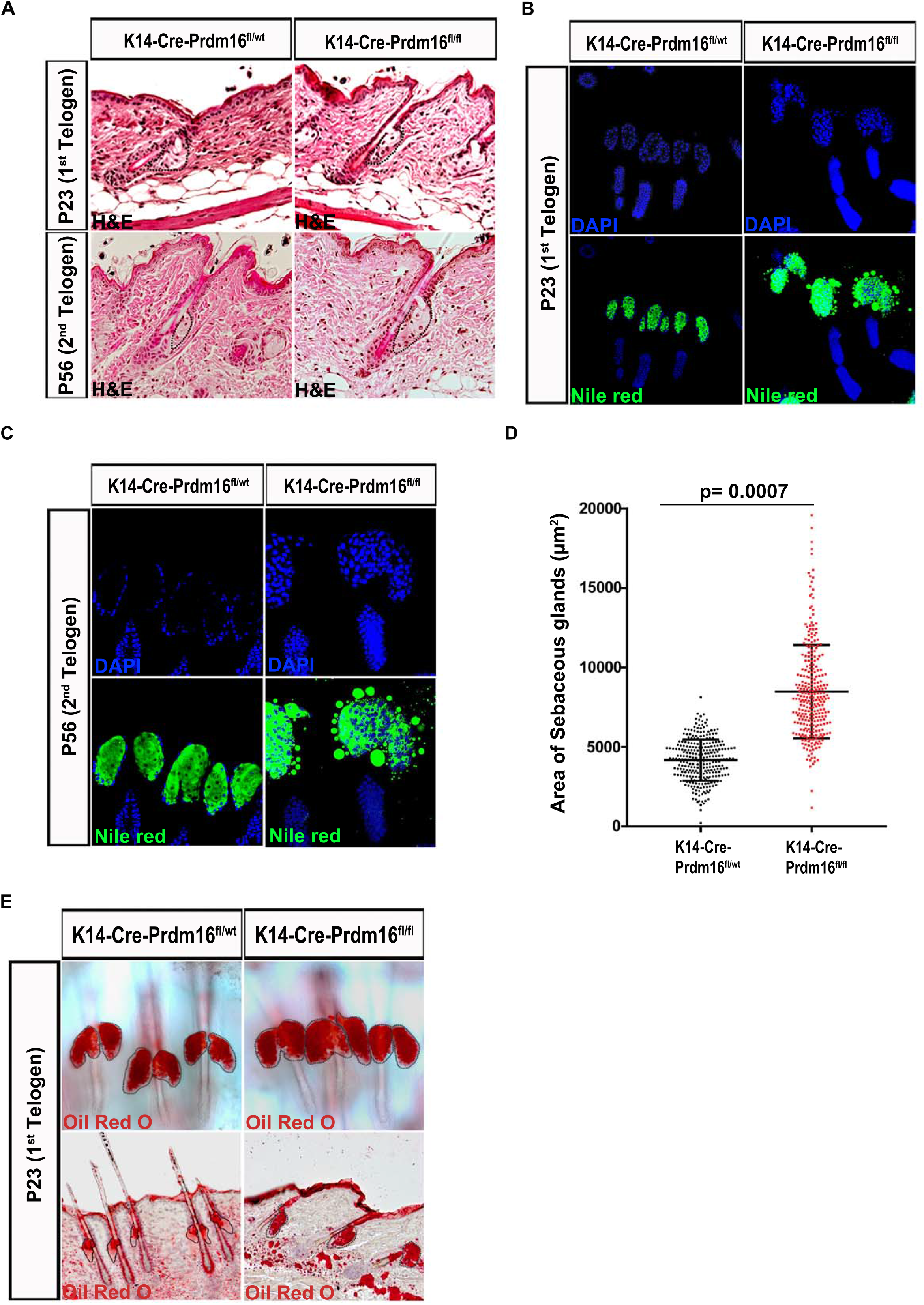
The conditional loss of PRDM16 in the skin causes an enlargement of the sebaceous glands and increased sebum production. **(A)** Back skin sections from P23 and P56 mice that were either WT (K14 Cre-Prdm16^fl/wt^; left) or deficient for PRDM16 in the epidermis (K14-Cre-Prdm16^fl/fl^; right) were stained with hematoxylin and eosin (H&E). The SGs, which appear enlarged in K14-Cre-Prdm16^fl/fl^ mice, are marked with a dotted line. **(B-C)** Back skin sections from (B) P23 (1^st^ telogen) and (C) P56 (2^nd^ telogen) mice that were either WT (K14 Cre-Prdm16^fl/wt^; left) or deficient for PRDM16 in the epidermis (K14-Cre-Prdm16^fl/fl^; right) were stained with Nile Red, which specifically labels sebum lipids. **(D)** Quantification of sebaceous gland sizes in back skin sections from P56 (2^nd^ telogen) mice that were either WT (K14 Cre-Prdm16^fl/wt^; left) or deficient for PRDM16 in the epidermis (K14-Cre-Prdm16^fl/fl^; right). Each dot represents a SG. For quantifications, four WT and four PRDM16 KO mice were used. The indicated p-value was calculated using an unpaired t-test (p=0.0007). **(E)** Tail epidermal sheets (upper row) and back skin sections (bottom row) from P23 (1^st^ telogen) mice that were either WT (K14 Cre-Prdm16^fl/wt^; left) or deficient for PRDM16 in the epidermis (K14-Cre-Prdm16^fl/fl^; right) were stained with Oil-Red-O.

### PRDM16 prevents excessive proliferation of sebocytes within the sebaceous gland

To examine if the observed SG enlargement and elevated sebum production in K14-Cre-*Prdm16^fl/fl^* mice was due to an increased number of differentiated sebocytes, we analyzed the expression of SCD1 and Adipophilin (ADRP), two markers of mature sebocytes (Miyazaki et al., 2001; Eisinger et al., 1993; Ostler et al., 2010), in back skin sections of 1^st^ and 2^nd^ telogen mice (Fig 4A, Fig 4 – figure supplement 1). These stainings clearly showed that the number of differentiated lipid-producing sebocytes was higher in K14-Cre-*Prdm16^fl/fl^* mice compared with control ones (Fig. 4A, Fig 4 – figure supplement). Next, to evaluate whether the elevated number of mature sebocytes in K14-Cre-*Prdm16^fl/fl^* mice could be caused by increased proliferation of JZ/SG SC and/or transiently amplifying sebocyte progenitors, we measured Ki67 expression levels. Remarkably, the loss of PRDM16 resulted in a dramatic increase in the number of proliferating cells within the JZ and the SG (Fig 4B-C). Moreover, while in control mice cell proliferation within the SG was restricted to the outer basal layer in K14-Cre-*Prdm16^fl/fl^* mice Ki67^+^ cells were also abundant within the inner part of the SGs where sebum-producing sebocytes accumulate (Fig 4B). Co-staining of SCD1 and Ki67 confirmed that, upon loss of PRDM16, differentiated sebocytes still proliferate (Fig 4D). These results suggest that seboctye proliferation and differentiation, two processes that are uncoupled during normal SG homeostasis (Horsley et al., 2006; Feldman et al., 2019), can occur simultaneously upon loss of PRDM16. It has been previously reported that C-MYC overexpression leads to enlarged SGs and a concomitant increase in both proliferation and sebocyte differentiation (Arnold and Watt, 2001; Waikel et al., 2001; Jensen and Watt, 2006; Horsley et al., 2006; Cottle et al., 2013). Interestingly, C-MYC was significantly overexpressed in the SGs of the K14-Cre-*Prdm16^fl/fl^* mice (Fig 4E), suggesting that PRDM16 controls SG homeostasis, at least partly, by repressing *c-myc* expression. Altogether, our work shows that PRDM16 is necessary for SG homeostasis and suggests that this transcription factor controls the balance between sebocyte proliferation and differentiation.

**Figure 4.**
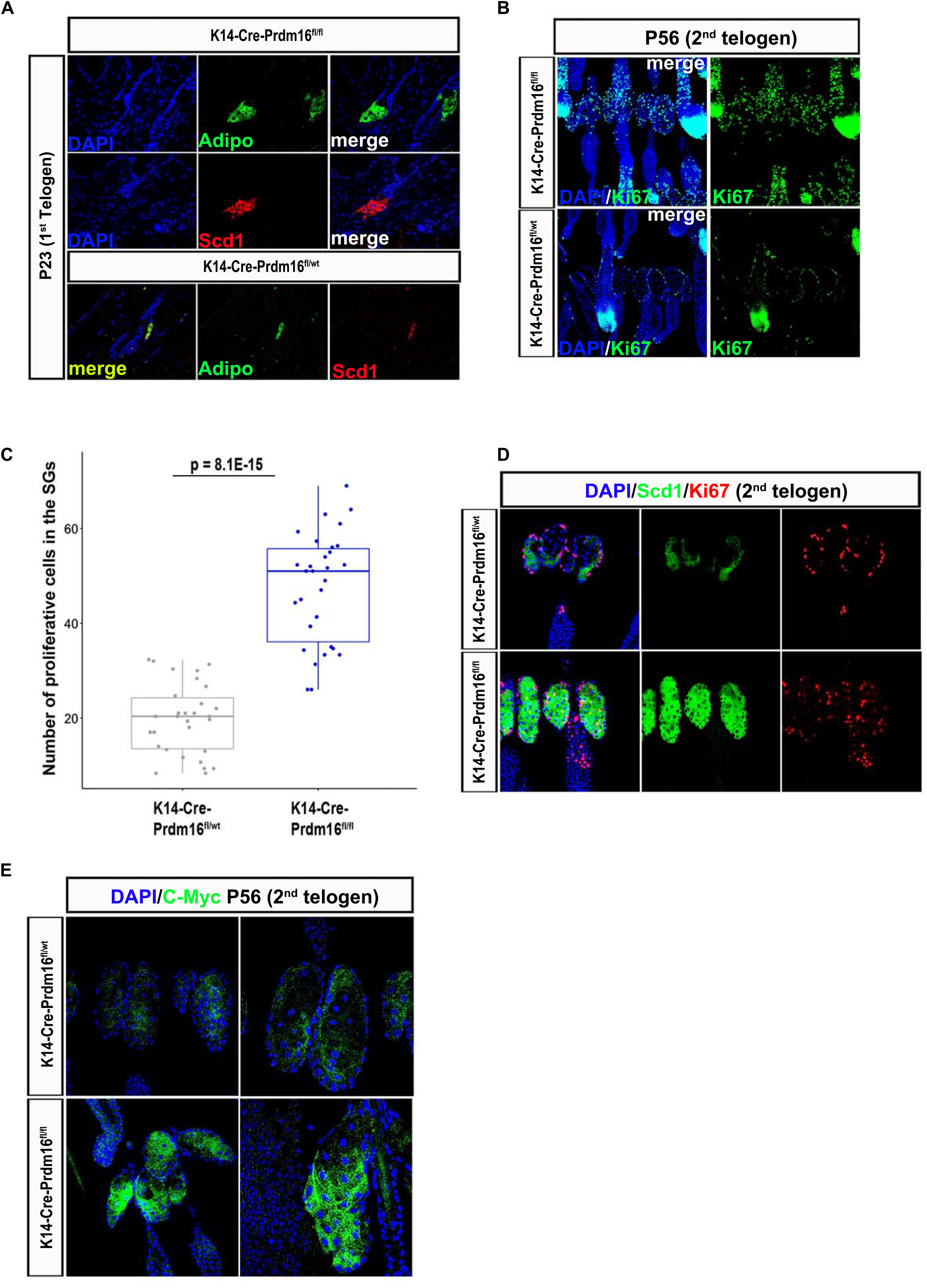
PRDM16 coordinates sebocyte proliferation and differentiation within the sebaceous gland. **(A)** Expression of SCD1 and Adipophilin (Adipo), two specific markers of differentiated sebocytes, was analyzed in back skin sections from P23 (1st telogen) mice that were either WT (K14 Cre-Prdm16^fl/wt^; bottom) or deficient for PRDM16 in the epidermis (K14-Cre-Prdm16^fl/fl^; top). **(B)** Expression of *Ki67*, a proliferation marker, specific was analyzed in back skin section from P56 (2^nd^ telogen) mice that were either WT (K14 Cre-Prdm16^fl/wt^; bottom) or deficient for PRDM16 in the epidermis (K14-Cre-Prdm16^fl/fl^; top). **(C)** Quantification of sebocyte proliferation was performed by counting the number of Ki67 positive cells within the SGs of P56 (2^nd^ telogen) mice that were either WT (K14 Cre-Prdm16^fl/wt^; bottom) or deficient for PRDM16 in the epidermis (K14-Cre-Prdm16^fl/fl^; top). Quantification was performed using three different Prdm16 KO mice and three different WT mice (60 sebaceous glands were obtained from each animal). The indicated p-value was calculated using an unpaired t-test. **(D-E)** Immunofluorescence was performed to analysed the expression of Ki67 (a known proliferative marker), SCD1 and C-Myc (specific markers of differentiated and differentiating sebocytes respectively) in tail skin whole mount sections from P56 (2^nd^ telogen) mice that were either WT (K14 Cre-Prdm16^fl/wt^; bottom) or deficient for PRDM16 in the epidermis (K14-Cre-Prdm16^fl/fl^; top).

## Discussion

We previously described a simple approach whereby H3K27me3 and RNA-seq profiles can be combined to functionally dissect cellular heterogeneity in various developmental contexts (Rehimi et al., 2016). Here we expanded the applicability of our epigenomic approach and used it to identify major regulators of poorly characterized SC populations. The epidermis is a major example of an adult epithelial tissue displaying spatially confined SC heterogeneity, whereby various SC populations participate in skin homeostasis and regeneration upon injury (Jaks et al., 2010; Schepeler et al., 2014). However, with the exception of the bulge HFSC, the additional SC populations residing with the IFE and the pilosebaceous unit remain poorly characterized at the molecular level and their key cell-identity regulators are only partially known. When applied to the bulk of SC present in the HF and the SG, our approach successfully identified previously reported non-bulge SC regulators. Most importantly, we also predicted important regulatory roles for several genes with no previously described functions in the HF or the SG SCs. For four of these potential SC regulators (e.g., *Prdm16, GJB6, MSX1 and DLK1*) we observed spatially restricted expression within non-bulge compartments of the pilosebaceous unit. Additional characterization of PRDM16, which was highly expressed in the JZ, uncovered a major role for this TF in SG homeostasis. Therefore, it is likely that many of the additional genes predicted as HF/SG regulators by our approach might play important functions in skin homeostasis and/or regeneration. More generally, our results indicate that our epigenomic approach can be a powerful tool to define the major cell identity regulators within adult stem cell populations, thus complementing current efforts to identify novel cell types both in the skin (Joost et al., 2016; Joost el al., 2018) as well as in other tissues using single-cell RNA-seq technology (e.g. *https://data.humancellatlas.org*).

*Prdm16* belongs to the PRDM family of TFs, whose members have major regulatory functions in several developmental and physiological contexts. Here we showed that *Prdm16*, which was previously described as a master regulator of brown fat cellular identity, plays an important role in SG homeostasis (Seale et al., 2008; Kajimura et al., 2010). Under homeostatic conditions, slow-cycling Lrig1+ SC located in basal layer of the JZ and the SG give rise to proliferative sebocyte progenitors (Kretzschmar et al., 2014; Page et al., 2013). As these progenitors are displaced towards the interior of the SG, they stop dividing, get fully differentiated and accumulate lipid droplets (Niemann and Horsley 2012). Although PRMD16 and LRIG1 display largely overlapping expression within the pilosebaceous unit, the loss of PRDM16 did not seem to affect the Lrig+ SC population residing within the JZ and SG basal layers. In contrast, in the absence of PRDM16, proliferation in the SG was no longer restricted to the basal layer but also frequently observed among lipid-producing sebocytes located in the SG interior. Consequently, *Prdm16^-/-^* mice displayed enlarged SGs and increased lipid production. Therefore, PRDM16 seems to be necessary to switch off proliferation in differentiating sebocytes. The regulatory networks that PRMD16 might control to ensure that proliferation ceases as sebocyte progenitors undergo terminal differentiation are still unknown. Nevertheless, the SG phenotypes observed in *Prdm16^-/-^* mice resemble those previously reported upon *Blimp1* deletion or *c-Myc* overexpression, respectively (Arnold and Watt, 2001; Horsley et al., 2006; Cottle et al., 2013; Kretzschmar et al., 2014). Furthermore, BLIMP1 and C-MYC negatively regulate each other expression in the SG (Horsley et al., 2006; Cottle et al., 2013; Kretzschmar et al., 2014). Therefore, we speculate that, once sebocyte progenitors start differentiating and producing sebum, PRDM16 might repress *c-Myc* and/or activate *Blimp1/Prdm1* in order to prevent their unrestricted proliferation. Further work will be required in order to identify the full repertoire of PRDM16 target genes during SG homeostasis.

In addition to increased sebocyte proliferation, the conditional loss of PRDM16 in the mouse epidermis resulted in excessive sebum production and secretion, a major pathogenic factor in the development of acne vulgaris (Moradi Tuchayi et al., 2015; Picardo et al., 2017; Zouboulis et al., 2004). Acne vulgaris is the most common disease of the pilosebaceous unit, but its pathomechanisms remain incompletely understood. A number of events are associated with the development of acne, which include not only increased sebum production, but also alterations in the quality of sebum lipids, inflammatory processes, dysregulation of the hormone microenvironment, interaction with neuropeptides, follicular hyperkeratinisation and the proliferation of propionibacterium acnes within the HF (Picardo et al., 2017; Moradi Tuchayi et al., 2015; Clayton et al., 2019). Interestingly, it was recently reported that GATA6, an important regulator of SG development and homeostasis (Oules et al., 2019; Oules et al., 2020), is downregulated in acne (Oules et al., 2020). Furthermore, GATA6 prevents hyperkeratinisation of the infundibulum and limits the lipid production and proliferation of sebocytes (Oules et al., 2020), which, as stated above, are two of the main pathological events associated with acne. We speculate that PRDM16 might constrain sebum production by activating *Gata6* in sebocyte progenitors. Therefore, analogously to GATA6 (Oules et al., 2020), PRDM16 might represent a relevant target for the future treatment of acne. However, additional work is still needed in order to identify the signalling pathways that control PRDM16 expression in the JZ and the SG and that could be therapeutically targeted in acne.

## Materials and Methods

### K14-Cre-Prdm16^fl/fl^ Mice

*Prdm16^lox/lox^* mutant mice with loxP sites flanking exon 9 of the *Prdm16* gene were ordered form Jackson lab (Stock No: 032160). To generate skin-specific Prdm16 knock-out mice, female *Prdm16^flox/flox^* mice were crossed with *Keratin-14 Cre* mice (K14-Cre; mixed Swiss Webster; C57Bl/6J; CBA/J; The Jackson Laboratory, Bar Harbor, ME), followed by backcrossing onto the C57Bl/6J background to generate compound heterozygous (*Prdm16^flox/+;Cre/+^)* mice. Male heterozygous *Prdm16^flox/+;Cre/+^* mice were subsequently mated with female *Prdm16^flox/flox^* (Lox) mice, generating *Prdm16^flox/flox;Cre/+^* mice. For litter expansion, male *Prdm16^flox/flox;Cre/+^* mice were bred with female Lox mice. All mice were on a fully backcrossed C57Bl/6J background, and Lox and *Prdm16^flox/flox;Cre/+^* mice were born in an expected ratio of 1:1. Genotyping was performed by PCR using genomic DNA isolated from a tail or ear clip, as described previously. The primers used for PCR were as follows. For K14 Cre genotyping, the forward primer: GTCCAATTTACTGACCGTACA and reverse primer: CTGTCACTTGGTCGTGGCAGC were used. For *Prdm16* flox genotyping, the mutant-specific forward primer: AGG AAC TTC ATC AGT CAG GTA CA, common reverse primer: TGC AGG GAG ATT GAC AAG TG and the WT-specific forward primer: ACC TTG AGG TTC TCG GTT AGA C was used (Cohen et al., 2014). Mice were maintained on a 12-h light/dark cycle with free access to water and either a standard chow diet (Purina 5008). Animals were housed and maintained according to FELASA guidelines in the animal facility of Centre for Pharmacology and Centre of Molecular Medicine Cologne, Cologne, Germany. All experiments were approved by local authorities. All animals were sacrificed by cervical dislocation and tissues (back skin and tail skin) were rapidly removed, either snap-frozen in liquid nitrogen or fixed in 4% PFA/1 x PBS and stored at −20 or −80 °C accordingly. All *in vivo* experimental procedures were approved by the Animal Care Research Committee of the University of Cologne (Germany).

### PSU SC isolation

Alpha6^+^/Sca1^-^ cells were isolated from back skin of P56 mice (2^nd^ telogen) by incubating skin pieces in 0.8% trypsin (Gibco) for 50 min at 37°C. After separating the epidermis from the underlying dermis, cells were passed through 70 µm and 45 µm cell strainers (BD Biosciences) and pelleted at 900 rpm for 7 minutes at 4 °C. Cells were resuspended in 250µl of FACS buffer (1 x PBS, 2.5% FBS or BSA and 2mM EDTA), afterward Alpha 6 conjugated antibody was added in each tube at concentration of 1:100. After vortex, samples were incubated at 4°C for 1 hr in the dark. The cells were washed twice with 1ml of FACS buffer and centrifuge at 900 rpm for 5 minutes at 4 °C. Cells were resuspended in FACS buffer. Cells were analysed in a BD FACS Canto II or sorted in either a BD FACSAria II or a BD FACSAria Fusion. Data were analysed using FlowJo software version 10. Expression of cell surface markers was analysed on live cells after exclusion of cell doublets and dead cells using 7AAD 1: 100 (eBioscience) or Fixable Viability Dye eFluor506 (eBioscience). The following antibody was used FITC-a6 Integrin (clone GoH3, eBioscience).

### ChIP-Seq

For each ChIP-seq experiment, 500,000 Alpha6^+^/Sca1^-^ cells isolated from back skin of P56 mice (2^nd^ telogen) were used. Alpha6^+^/Sca1^-^ cells were briefly homogenized in DMEM media with 10% FBS serum/1M sodium butyrate and cross-linked with 1% formaldehyde at room temperature with rotation for 15 min. Cross-linking was quenched with 0.125 M glycine, and HFSC were rinsed with cold 1 X PBS, re-suspended in 500 µl of sonication buffer (10 mM Tris, 100 mM NaCl, 1 mM EDTA, 0.5 mM EGTA, 0.1% Na-deoxycho-late, and 0.5% N-lauroylsarcosine) with 1% SDS, and incubated at 4C for 10 min. Chromatin was sonicated to generate DNA fragments of 200–500 bp. Sonicated chromatin was incubated overnight with 5 µg of antibodies against H3K27me3 (39536; Active Motif) or H3K4me3 (39159; Active Motif; Rada-Iglesias et al., 2011; Rehimi et al., 2016), followed by 4–6 hr incubation with Protein G Dynabeads (Life Technologies). Beads were washed four times with 1 mL of cold radioimmunoprecipitation assay buffer (RIPA) washing buffer (50 mM HEPES, 500 mM LiCl, 1 mM EDTA, 1% NP-40 and 0.7% Na-deoxycholate) and once with Tris/EDTA buffer (TE) containing 50mM NaCl. Immunoprecipitated chromatin was eluted from the beads by adding elution buffer (50 mM Tris, 10 mM EDTA, and 1% SDS) and vortexing at 65C for 15 min. Crosslinking was reversed by incubating the eluted samples at 65C overnight. Samples were then diluted (1:1) with TE and sequentially treated with RNase A (0.2mg/mL) and proteinase K (0.2mg/mL). DNA was purified using phenol:chloroform, followed by ethanol precipitation, and re-suspended in water. DNA libraries from H3K27me3 ChIP, H3K4me3 ChIP, and corresponding input DNA samples were generated with the TruSeq kit (Illumina) as single reads. Sequences were mapped with Burrows-Wheeler Aligner (BWA) (Li and Durbin 2009) to the mouse mm10 genome assembly. The resulting Binary Alignment/Map (BAM) files for the H3K27me3 ChIP-seq data were analyzed with MACS (Zhang et al., 2008) using the following settings: –broad–gsize 8e8 -q 0.1 -m 5 50–fix-bimodal–ext sizes (Zhang et al., 2008). Among the identified peaks, only those with fold-enrichments ≥3 were considered for downstream analyses. Finally, peaks located ≤2 kb from each other was clustered into single genomic intervals using Galaxy tools. The generated ChIP-seq datasets in Alpha6^+^/Sca1^-^ cells are accessible through GEO: (<u>GSE169647</u>).

### RNA-Seq

Approximately ∼10^5^ Alpha6^+^/Sca1^-^ cells from back skin of P56 mice (2^nd^ telogen) were isolated and collected in 1.5 ml Eppendorf from 56 days old mice (P56: 2nd telogen). Accumulated cells were used to isolate total RNA using the RNeasy Plus Micro Kit after which the RNA quality was determined using an Agilent 2200 Tape Station. Libraries were prepared and sequenced as previously described as three independent biological replicates (Rehimi et al., 2016). The generated RNA-seq datasets in Alpha6^+^/Sca1^-^ cells are accessible through GEO: (<u>GSE169648</u>).

RNA-seq data from bulge HFSC was previously generated as three biological replicates (Chacon-Martinez et al., 2017) and was retrieved from GEO (GSE76779).

### Differential Functional Heterogeneity (dFH) Score

The dFH score was computed as follows:

#### dFH_g_= (H3K27me3B_g_+1) x log_2_(DE_g__FoldChange) x -log(DE_g__qValue)

- *H3K27me3B_g_* is the breadth of the H2K27me3 mark in Alpha6^+^/Sca1^-^ cells at the gene *g*. We added 1 to the H3K27me3 breadth to avoid mathematical issues with zeros.
- *log*_2_(*DE*_*g*_*_FoldChange*) is the log_2_ gene expression fold-change of the gene *g* calculated through differential gene expression comparing Alpha6^+^/Sca1^-^ cells against bulge HFSC using DESeq2 (Love et al., 2014).
- *-log(DE_g__qValue)* represents the significance of the differential expression (measured as –log(q-value)) of the gene *g* when comparing Alpha6^+^/Sca1^-^ cells against bulge HFSC using DESeq2 (Love et al., 2014).

All mouse genes and their corresponding dFH scores are listed in Data S1.

### Haematoxylin and Eosin (H&E) staining

Fresh back skin samples from similar sites obtained from age and gender-matched control and *Prdm16* Knockout mice were fixed in 4% PFA/PBS at RT for 30 minutes. Subsequently, back skin samples were paraffin-embedded for sectioning (8µm thick) and stained with hematoxylin (Sigma) at RT for 3 minutes. Counterstaining was achieved by rinsing with eosin (Sigma) for 30 seconds, followed by dehydration through sequential washing with 95% ethanol and 100% ethanol. Slides were mounted with Aqua Poly/Mount (18606-20, Polysciences, Inc).

### Oil Red O Staining

Back skin sections between 7 and 12 µm thick were cut on a cryostat (LEICA CM 3050S) and attached to Super Frost Plus slides (Menzel-Glaser; Thermo Scientific), air dried for up to 90 minutes at room temperature (RT), and then stained with Oil Red O dye (O0625-25G, Sigma) to detect the presence of lipids. Cryo-sections and tail skin whole mounts were washed three times in PBS and fixed in 4% Paraformaldehyde (Sigma) in PBS for 30 mints at RT. Next, samples were incubated for 10 minutes in 60% isopropanol (Sigma) and then stained for 30 minutes with Oil Red O solution. Samples were then briefly rinsed in 60% isopropanol, washed thoroughly in water, counterstained in Mayers Hematoxylin solution (Fluka) and mounted under coverslips in Aqua Poly/Mount (18606-20, Polysciences, Inc).

### Nile Red staining

Fresh tail whole mounts samples from similar sites obtained in age and gender-matched control and *Prdm16* Knockout mice were fixed in 4% PFA/PBS for 30 minutes at RT. Samples were washed once in PBS for 10 minutes at RT. Next, samples were incubated in 1µl Nile Red (N3013, Sigma)/1ml PBS for 30 minutes at RT and nuclei were stained with DAPI (Sigma) at RT for 15 minutes. Slides were mounted with Aqua Poly/Mount (18606-20, Polysciences, Inc).

### Immunofluorescence and immunohistochemistry

Tissue samples were fixed in 4% Paraformaldehyde/PBS for 1hr at RT, dehydrated, and embedded for cryo- and paraffin-sections. Sections (7-12µm) were deparaffinized, rehydrated, permeabilized in 0.5% Triton X-100/1xPBS for 10 minutes and blocked with 2% BSA/PBS/0.5% Triton X-100 for 30 minutes at RT. Sections were stained with primary antibodies in 0.5% Triton X-100, 1% BSA, PBS overnight at 4C. Secondary, fluorescent antibodies were used to detect primary antibody binding, and nuclei were visualized with DAPI. Slides were mounted with Aqua Poly/Mount (18606-20, Polysciences, Inc). The following primary antibodies were used: PRDM16 (PA520872, Thermo Fisher Scientific), MSX1 (ab174207, Abcam), DLK1 (3A10, Thermo Fisher Scientific), GJB6 (ab200866, Abcam), SCD1 (SC14719, Santa Cruz Biotechnology), LRIG1 (AF3688, R&D systems) and Adipophilin (20R-AP002, Fitzgerald Industries International). Secondary antibodies (anti-mouse AlexaFluor 488 and anti-rabbit Alexa Fluor 568) were purchased from Life Technologies.

### Imaging and Fiji (Image J) Analysis of Skin Samples

Imaging of immunofluorescence and histological stainings were performed on Olympus BX 53 or Olympus Fluoview FV 1000 with their corresponding softwares, respectively. To quantifying the number of proliferative cells or the size of the SGs in *Prdm16* knockout and WT mice, images of tail whole mounts obtained as described above were analyzed using Fiji (ImageJ-win32) software. The *cell count* function in plugins was used to count the proliferative cells and the *free hand* function in toolbar was used to draw the outlines around the regions of interest. After this, the total numbers of proliferative cells were counted and the size of SGs was measured per square micrometer (µm^2^).

---

**Figure 1 – figure supplement 1.**
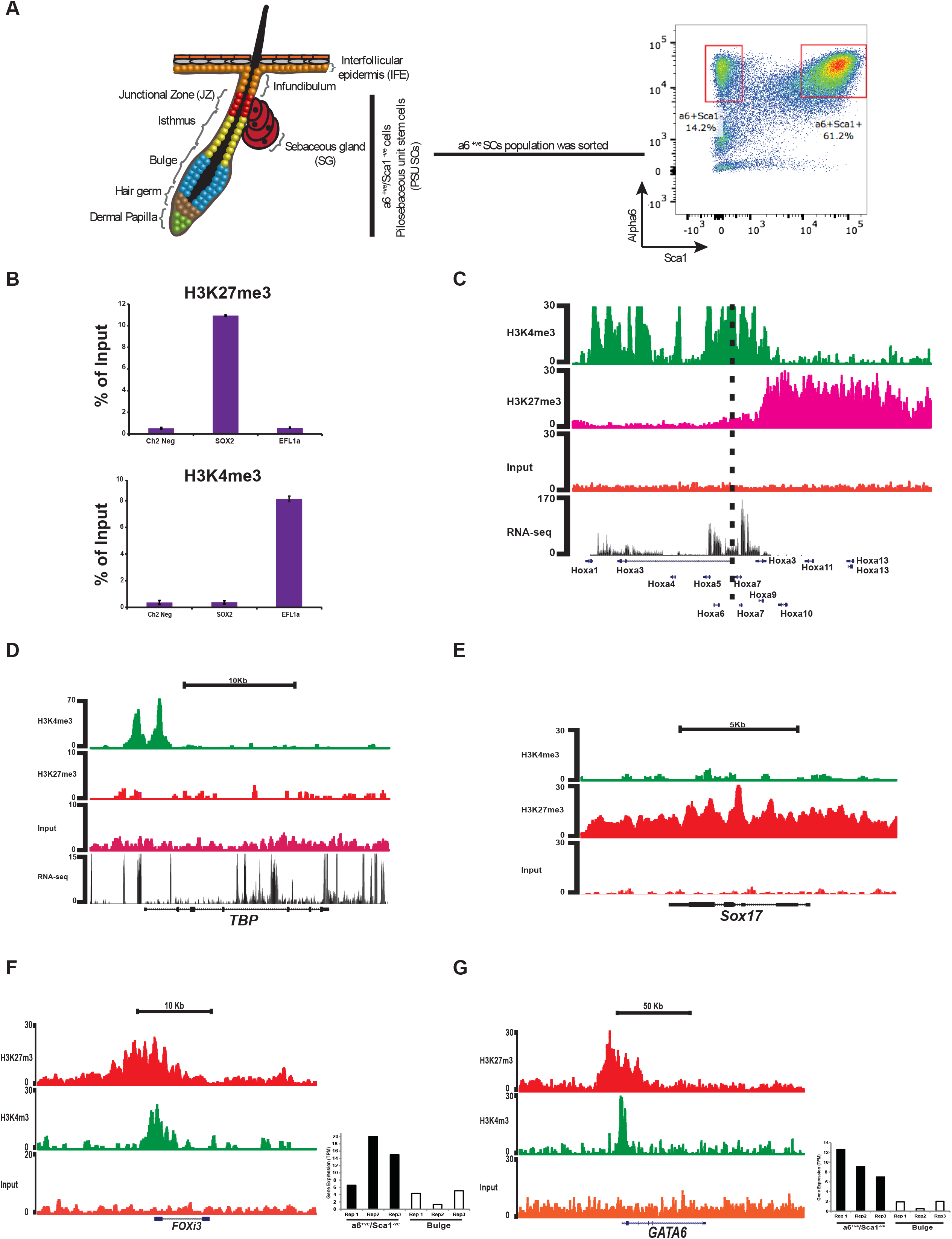
Epigenomic analysis of sorted epidermal stem cells. **(A)** Left: Schematic diagram showing different epidermal compartments and the location of α6^+ve^/Sca1^−ve^ PSU SC. Right: representative FACS plot of freshly isolated epidermal cells stained with the indicated antibodies. The α6^+ve^/Sca1^−ve^ and α6^+ve^/Sca1^+ve^ populations are highlighted. **(B)** H3K27me3 and H3K4me3 levels at the indicated regions were measured by ChIP-qPCR analysis using α6^+ve^/Sca1^−ve^ cells isolated from P56 adult mice skin. Chr2 Neg corresponds to an intergenic region in mice chromosome 2 and serves as a H3K27me3 negative control; the promoter of *Sox2*, which is inactive in α6^+ve^/Sca1^−ve^ HFSC, is expected to be marked by H3K27me3 but not H3K4me3; the promoter of *Eef1a*, a housekeeping gene active in α6^+ve^/Sca1^−ve^ PSU SC, is expected to be marked by H3K4me3 but not by H3K27me3. **(C)** ChIP-seq (H3K4me3 and H3K27me3) and RNA seq profiles generated from α6^+ve^/Sca1^-ve^ stem cells around the HOX locus. The HOXA cluster can be clearly divided in active (high RNA-seq and high H3K4me3 levels) and inactive (low RNA-seq and high H3K27me3) domains, which illustrates the quality of the ChIP-seq data generated from sorted α6^+ve^/Sca1^-ve^ PSU SC. **(D-E)** ChIP-seq and RNA-seq profiles from sorted α6^+ve^/Sca1^-ve^ PSU SC around a housekeeping gene (*Tbp*) and a developmental gene not expressed in the epidermis (*Sox17*). **(F and G)** ChIP-seq (H3K27me3 in red, H3K4me3 in green and input in orange) profiles and RNA-seq values (bar plots) are shown for *Gata6* and *Foxi3*, two genes recently described as non-bulge SC regulators (Wang et al., 2017; Shirokova et al., 2016; Donati et al., 2017).

**Figure 2-figure supplement 1.**
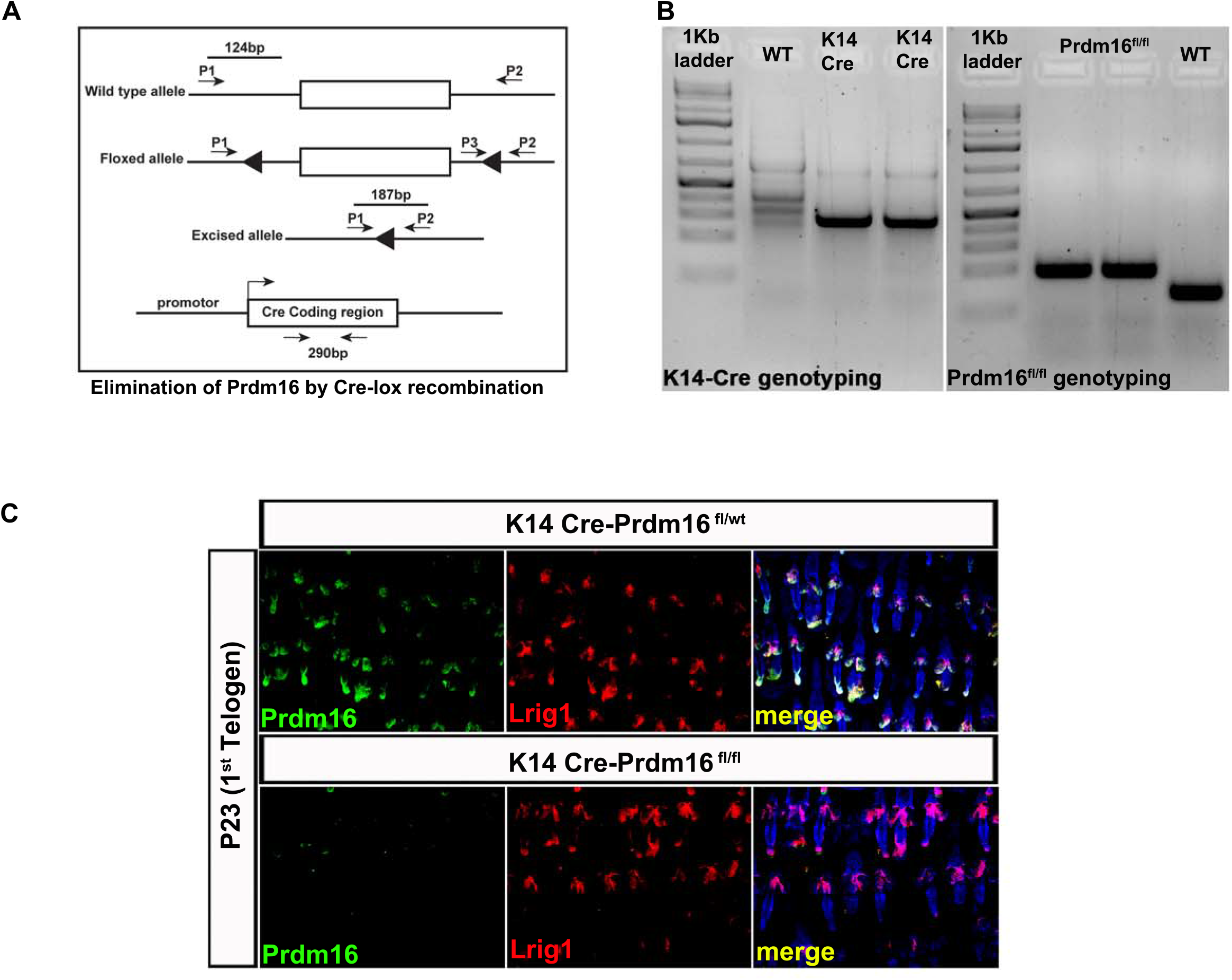
Epidermal-specific Prdm16 Knock-out mouse model. **(A)** Schematic diagram of the PCR strategy used to confirm conditionally loss of Prdm16 in the Alpha6^+^/Sca1^-^ PSU SC. Three primers P1 (Mutant-specific forward), P2 (Wild type-specific forward) and P3 (Common) were used to amplify exon 9 of the Prdm16 gene. **(B)** PCR analysis of DNA from 21 days old K-Cre-Prdm16 ^flox/flox^ mouse isolated from tail biopsy confirmed the deletion of Prdm16 gene in K14-Cre mice. **(C)** Double immunofluorescence for PRDM16 and LRIG1 in back skin sections from P23 (1^st^ telogen) mice that were either WT (K14 Cre-Prdm16^fl/wt^; upper row) or deficient for PRDM16 in the epidermis (K14-Cre-Prdm16^fl/fl^).

**Figure 3 – figure supplement 1.**
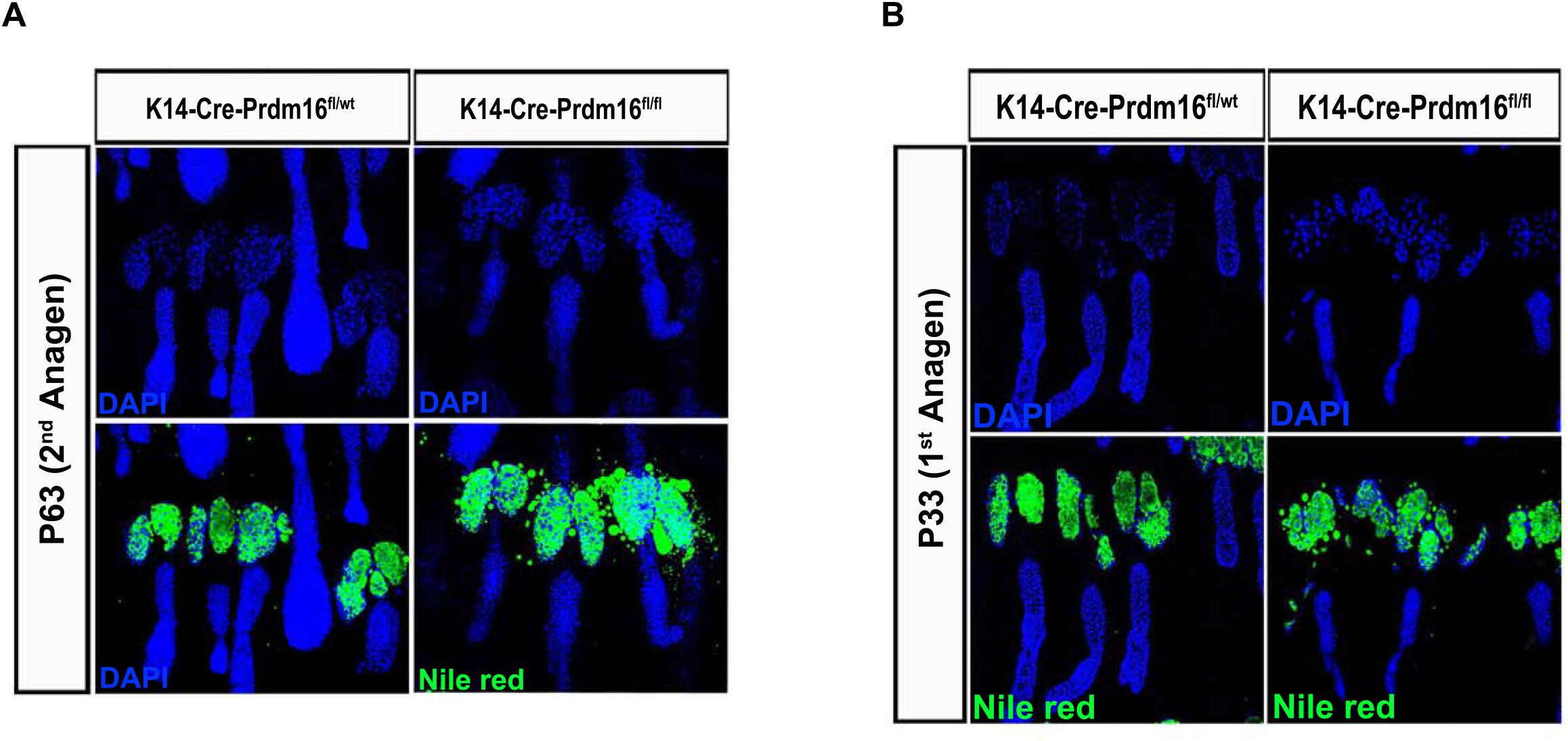
PRDM16 deficient mice display enlarged sebaceous glands and increased sebum production. **(A-B)** Back skin sections from (A) P63 (2^nd^ anagen) and (B) P33 (1^st^ anagen) mice that were either WT (K14 Cre-Prdm16^fl/wt^; left) or deficient for PRDM16 in the epidermis (K14-Cre-Prdm16^fl/fl^; right) were stained with Nile Red.

**Figure 4 – figure supplement 1.**
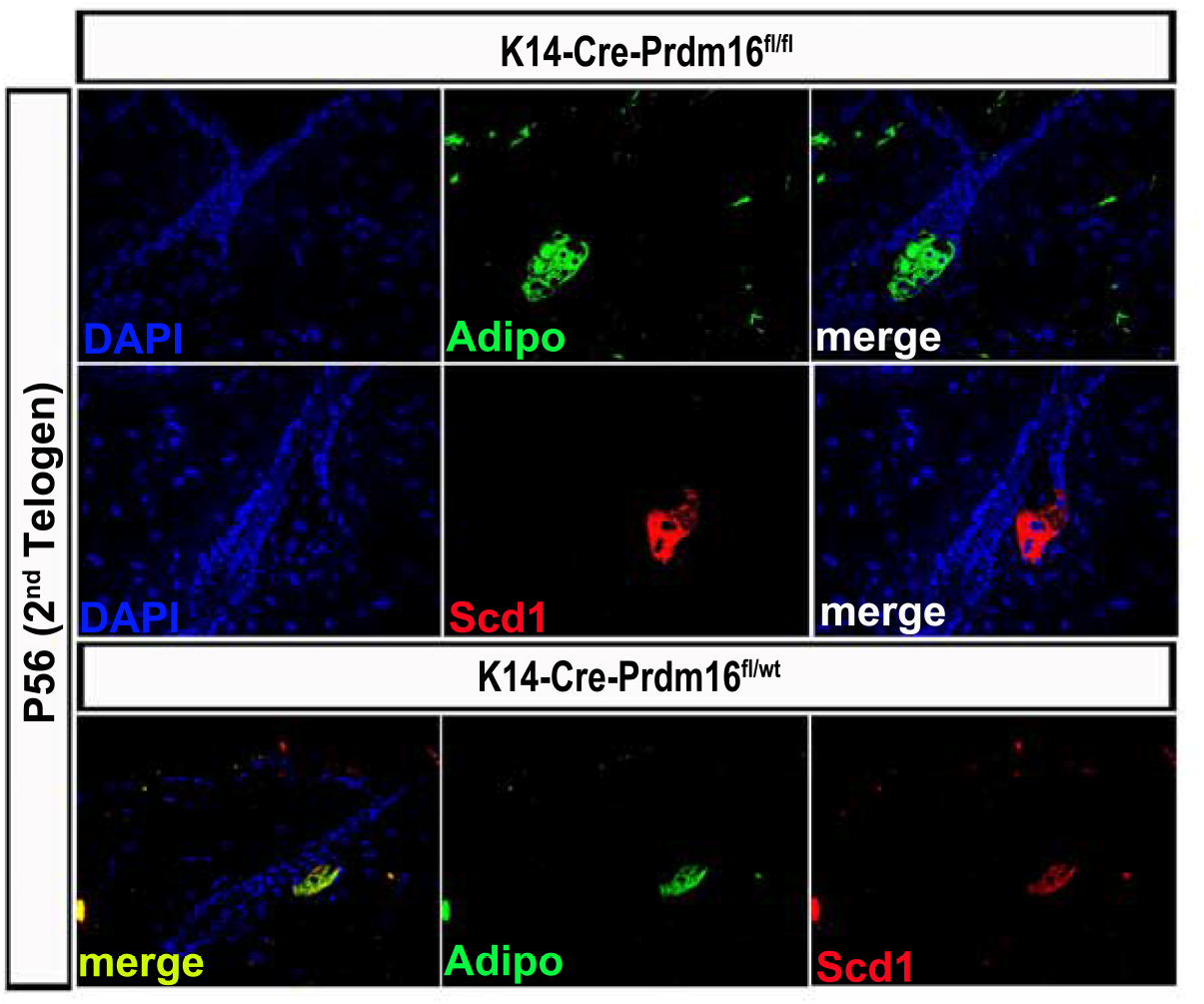
PRDM16 deficient mice display higher number of differentiated lipid-producing sebocytes. Expression of SCD1 and Adipophilin (Adipo), two specific markers of differentiated sebocytes, was analyzed in back skin sections from P23 (1st telogen) mice that were either WT (K14 Cre-Prdm16^fl/wt^; bottom) or deficient for PRDM16 in the epidermis (K14-Cre-Prdm16^fl/fl^; top).

## Notes

### Competing Interest Statement

The authors have declared no competing interest.

